# The gut commensal *Akkermansia muciniphila* ameliorates dysbiosis-mediated transplant arterial injury

**DOI:** 10.64898/2026.08.26.747435

**Authors:** Jenice M. Dumlao, Kevin Rey, Paige McCallum, Ethan Wheatley, Winnie Enns, Colten R. Hodak, Lauren E. Davey, Jonathan C. Choy

## Abstract

**Background:** Transplant arterial injury is an underlying feature of acute organ transplant rejection and is a main cause of late heart transplant failure. The role of the gut microbiota, and especially specific microbial components of this community, in controlling immune responses that cause this aspect of rejection is poorly understood.

**Methods:** We utilized a murine aortic interposition model of transplant arterial injury to investigate the role of the gut commensal bacteria, *Akkermansia muciniphila*, in controlling immune responses in transplant arteries.

**Results:** Early life treatment of female mice with broad spectrum antibiotics, which delayed colonization of the intestinal tract with bacteria until after weaning, led to the development of dysbiosis in adults that was characterized by the absence of *A. muciniphila*. This was related to an elevation in systemic levels of CCL2 and a reduction in the immunomodulatory short-chain fatty acid, propionate. When transplant arterial injury was examined, there was more arterial injury indicative of acute rejection and increased intimal thickening reflective of transplant arteriosclerosis in grafts from dysbiotic mice compared to controls. Dysbiosis also increased macrophage accumulation early after transplantation in dysbiotic mice. Notably, restoring *A. muciniphila* in the gut microbiota of dysbiotic mice through voluntary oral administration in infants ameliorated macrophage-mediated transplant arterial injury.

**Conclusions:** *A. muciniphila* is an immunomodulatory component of the gut microbiota that protects against vascular injury and pathology in organ transplantation.

## Introduction

The gut microbiota is the community of viruses, bacteria and fungi that inhabit the intestinal tract and plays a vital role in many aspects of health and disease. This microbial population provides microbial signals and metabolites to epithelial and immune cells in the intestine^1^. In this way, it tailors local and systemic immune activation to environmental exposures that affect it^2^. One of the main ways that the gut microbiota activates or regulates immune responses in non-intestinal tissues is by ;roducing metabolites that act on the intestinal epithelium to enhance its barrier properties or that can cross the intestinal epithelium into circulation, where they reach sites distal to the gut^3–6^. A reduction in intestinal barrier function leads to the systemic dissemination of microbial signals where they can enhance the activation status of innate immune cells to promote immune activation ^7–9^. Some metabolites, such as short chain fatty acids (SCFAs), also directly permeate through the intestinal epithelium into the blood stream where they disseminate systemically and regulate innate immune cell activation by stimulating G-protein coupled receptors and inhibiting histone deacetylases^10–12^. Dysbiosis, which is the imbalance or alteration in the composition of the gut microbiota that has detrimental effects to the host, can be induced by environmental factors such as early-life antibiotic use^13,14^. It is well-established that dysbiosis caused by early life antibiotic exposure leads to a reduction in many pro-health microbial features, which contributes to the subsequent development of immune-mediated diseases such as asthma, cardiovascular disease and inflammatory bowel disease^13,15–17^.

*Akkermansia muciniphila* is a commensal gut bacterium that comprises 1-4% of the total fecal gut microbial population^18^. It is present across many mammalian species, including humans and rodents, where it is the only bacterium that uses carbohydrates from mucin as its sole energy source^18,19^. This bacterium is considered a pro-health microbe because its abundance is negatively associated with several cardiovascular and metabolic diseases^20,21^. Specifically, experimental studies show that this bacterium protects against atherosclerosis, metabolic syndrome, and type 2 diabetes in mouse models by maintaining intestinal barrier properties that prevents the dissemination of bacterial components and signals into the circulation ^20–23^. Notably, *A. muciniphila* is being examined in clinical trials for the prevention of metabolic changes related to type 2 diabetes and early findings suggest that this bacterium improves metabolic health^24,25^. *A. muciniphila* stimulates the expression of tight junctional proteins by inducing TLR signaling^26^. This bacterium also produces the SCFAs acetate and propionate, which have barrier-inducing effects on intestinal epithelial cells as well as immunoregulatory properties on myeloid cells ^12,27–29^.

The gut microbiota functions as an inter-dependent population, with its physiological effects reliant on its taxonomic and/or metagenomic composition^6^. This composition is defined early in life when the intestinal tract begins to be colonized with microbes immediately after birth^2^. It is known that colonization of the gut microbiota in infancy is important for proper development of immune and metabolic homeostasis, and is controlled by mode of delivery, breastmilk and early exposure to environmental factors like diet and antibiotics^30^. Disruptions of the gut microbiota during infancy can have profound effects on immune and metabolic disorders that develop later in adolescence and adulthood, with antibiotic use in infancy being associated with increased likelihood of developing obesity and atopic dermatitis in adolescents^31,32^. Amoxicillin exposure to piglets also transiently alters the gut microbial composition, which leads to increased IFNγ production in peripheral blood mononuclear cells and increased neutrophil activity during inflammatory challenge later in life^33^. Colonization during the weaning period of development is also recognized as an essential component of the healthy gut microbiota. Alterations in microbial exposure during this early period of life reduces intestinal barrier function through effects on the proliferation of intestinal stem cells^34,35^. As such, it is important to understand how dysbiosis caused by antibiotic exposure and altered colonization during weaning affects the immunological properties of the gut microbiota.

The gut microbiota is important in organ transplantation^14,36–38^. Low microbial diversity and a reduction in SCFA-producing bacterial populations have been associated with increased post-operative infections and mortality in kidney, liver, and heart transplant recipients^39–42^. Some pre-clinical studies show that eliminating the microbiota before skin and lung transplantation reduces type I interferon signalling in antigen-presenting cells, thereby reducing alloimmunity and transplant rejection^43^. There are also immunoregulatory features of the gut microbiota because it promotes the production of regulatory IL-10 B-cells that induce tolerance against skin and cardiac allografts in mice^44,45^. Specific bacterial species can also be delivered to graft recipients to inhibit rejection, with oral gavage of *Alistipes onderdonkii* able to prolong skin allograft survival^37^ and *Bifidobacterium pseudolongum* able to improve cardiac allograft outcomes^37^.

Immune-mediated injury of transplant arteries is a component of acute and chronic heart transplant rejection^46^. Arterial injury during acute rejection is characterized by endothelialitis, leukocyte infiltration into the adventitia and media, and resultant injury to vascular endothelial and smooth muscle cells^46^. Early endothelial injury and persistent immunological targeting of arteries leads to intimal thickening, which is reflective of transplant arteriosclerosis (TA) that is a leading cause of late heart transplant failure^46^. We previously showed that delaying the colonization of the gut microbiota until after weaning by giving infant mice broad spectrum antibiotics until 3 weeks of age results in dysbiosis characterized by a loss of *A. muciniphila*^14^. This dysbiosis caused an exacerbation of myeloid cell-mediated transplant arterial injury in female graft recipients ^14^. In this current study, we investigate the influence of *A. muciniphila* on transplant arterial injury by colonizing dysbiotic mice with this bacterium. Our results show that dysbiosis of the female murine gut microbiota leads to a reduction in the thickness of the colonic mucosal layer and in intestinal levels of the SCFA propionate, both of which are features of reduced barrier function. There is also increased systemic levels of CCL2 in dysbiotic mice that lack *A. muciniphila,* suggestive of enhanced systemic inflammation. Dysbiosis led to exacerbation of transplant arterial injury characterized by increased medial injury mediated by early macrophage infiltration into transplanted allografts and intimal thickening at later time-points. Notably, restorative colonization of *A. muciniphila* in the gut by voluntary oral feeding of infants completely abrogates the enhanced transplant arterial injury cause by dysbiosis. These findings establish *A. muciniphila* as an anti-inflammatory commensal microbe that protects against transplant arterial injury in female graft recipients.

## Methods

### Animals

Balb/c and C57Bl/6 mice were purchased from Jackson laboratories and bred in-house. Colonization of the gut microbiota was delayed in C57Bl/6 mice by administering an antibiotic cocktail containing ampicillin (0.5 g/L), vancomycin (0.25 g/L), neomycin sulfate (0.5 g/L), and metronidazole (0.5 g/L) in the drinking water until pups were 3 weeks old as previously described^14,36^. Antibiotic water was administered ad libidum to pregnant mothers and pups. Mice were maintained on regular water after 3 weeks of age. All study protocols were reviewed and approved by the Simon Fraser University Animal Care Committee.

### Cytokine Array

100µL of whole blood was collected from mice at 12 weeks of age. Whole blood was centrifuged at 1000g for 10 mins at 4°C, and serum collected. Serum samples were diluted 1:2 and cytokine and chemokine levels analyzed using a bead-based mouse cytokine/chemokine 32-plex ELISA (MD32; Eve Technologies, Calgary, AB).

### Short-chain fatty acid analysis

Liquid chromatography–mass spectrometry was performed on fecal and serum samples at The Metabolomics Innovation Centre (University of Alberta, Edmonton, Canada) to quantify acetate, butyrate, and propionate.

### Metagenomic shotgun sequencing and bacterial composition analysis

Whole-genome shotgun sequencing was performed at the Michael Smith Genome Sciences Center (Vancouver, British Columbia, Canada) on the Illumina HiSeq Platform and analyzed using the bioBakery pipeline^47^. Briefly, raw reads were filtered for quality and contaminating mouse DNA using KneadData^47^. The composition of bacterial species was determined using MetaPhlAn4^48^ and significant differentially abundance species were determined using MaAsLin3^49^.

### Culture of *Akkermansia muciniphila*

*A. muciniphila (*MucT/BAA-835) was obtained from ATCC. Cultures were grown in a previously described synthetic medium^50,51^. Cultures were incubated in an anaerobic chamber (Coy Laboratory; 5% H₂, 5% CO₂, 90% N₂) until they reached stationary phase. Cells were harvested by centrifugation, washed once with PBS, then resuspended to approximately 10¹⁰ CFU/mL in sterile anaerobic PBS containing 20% glycerol.

### Colonization of the gut with *A. muciniphila*

Within the first week after weaning from antibiotics, *A. muciniphila* was administered to mice through voluntary feeding with a sweetened gelatin capsule as described previously^51^. Briefly, mice were fasted for 3 hours and then trained to eat a 0.25g sucralose-sweetened control gelatin capsule containing no bacteria. 1-2 days after training, mice were then administered a gelatin capsule containing 10^9^ CFU of *A. muciniphila*. Stool samples were collected up until 56 days after probiotic feeding (11 weeks of age). The presence and absence of endogenous mouse *A. muciniphila* (mAKK) was determined by PCR as previously described^51^. mAKK abundance relative to pan-16S and absolute quantification of exogenous human *A. muciniphila* (hAKK) was measured by quantitative PCR.

### Aortic Interposition Grafting

Artery transplants were performed as previously described^14,36^. Briefly, 8-to-12 week old C57Bl/6 mice received a section of infrarenal aorta from sex-matched Balb/c donors that was interposed into the infrarenal aorta of recipients. Artery segments from C57Bl/6 mice were used as syngraft controls to show no rejection.

### Histology and Immunohistochemistry

Artery grafts were harvested and perfusion fixed immediately in 4% paraformaldehyde. Proximal colon samples were harvested and fixed in methaCarnoy’s solution (methanol, 30% chloroform, 10% acetic acid). All tissue samples were snap frozen in optimal cutting temperature media (OCT, Thermofisher) and sectioned into 8µm slices prior to histological staining.

Cross sections of arterial grafts were H&E stained and medial thickness, intimal thickness and % luminal narrowing quantified as previously described^14^. Immunohistochemistry was performed to quantify immune cell infiltration and vascular injury in artery sections using antibodies against myeloperoxidase (MPO, Abcam), Mac-3 (BDBiosciences), CD4 (4SM95, eBiosciences), CD8 (4SM15, eBiosciences) as previously described (citations).

### Immunofluorescence

Proximal colon cross sections were rehydrated in PBS, then stained with UEA-1-FITC (GeneTex, GTX01512) for 30 minutes to visualize the colonic mucosa. Images were acquired on a Zeiss LSM 800 Airyscan with 40X/1.1 W objective lens and processed with the Zeiss Zen black software.

### Statistics

To determined differences between multiple groups, a Kruskal Wallis and Dunn’s post-hoc test was performed and a Mann Whitney U Test was performed for comparison between two groups. Statistically significant differences were set at p<0.05.

## Results

### Dysbiosis leads to reduced short-chain fatty acid production and increased systemic inflammation

We showed previously that dysbiosis of the female murine gut microbiota caused by early-life treatment with antibiotics exacerbates immuneimediated injury of transplant arteries that is associated with increased myeloid cell accumulation shortly after transplantation ^36^. Given the known role of the gut microbiota in training the innate immune system, we examined whether there are alterations to systemic inflammation that may be caused by this dysbiosis. Serum levels of 32 cytokines and chemokines were measured before transplantation to investigate baseline systemic inflammation. Notably, the monocyte chemoattractant CCL2 (MCP-1) was markedly and significantly elevated in dysbiotic female mice compared to controls (Figure 1A), suggesting a heightened set-point for myeloid cell activation.

**Figure 1.**
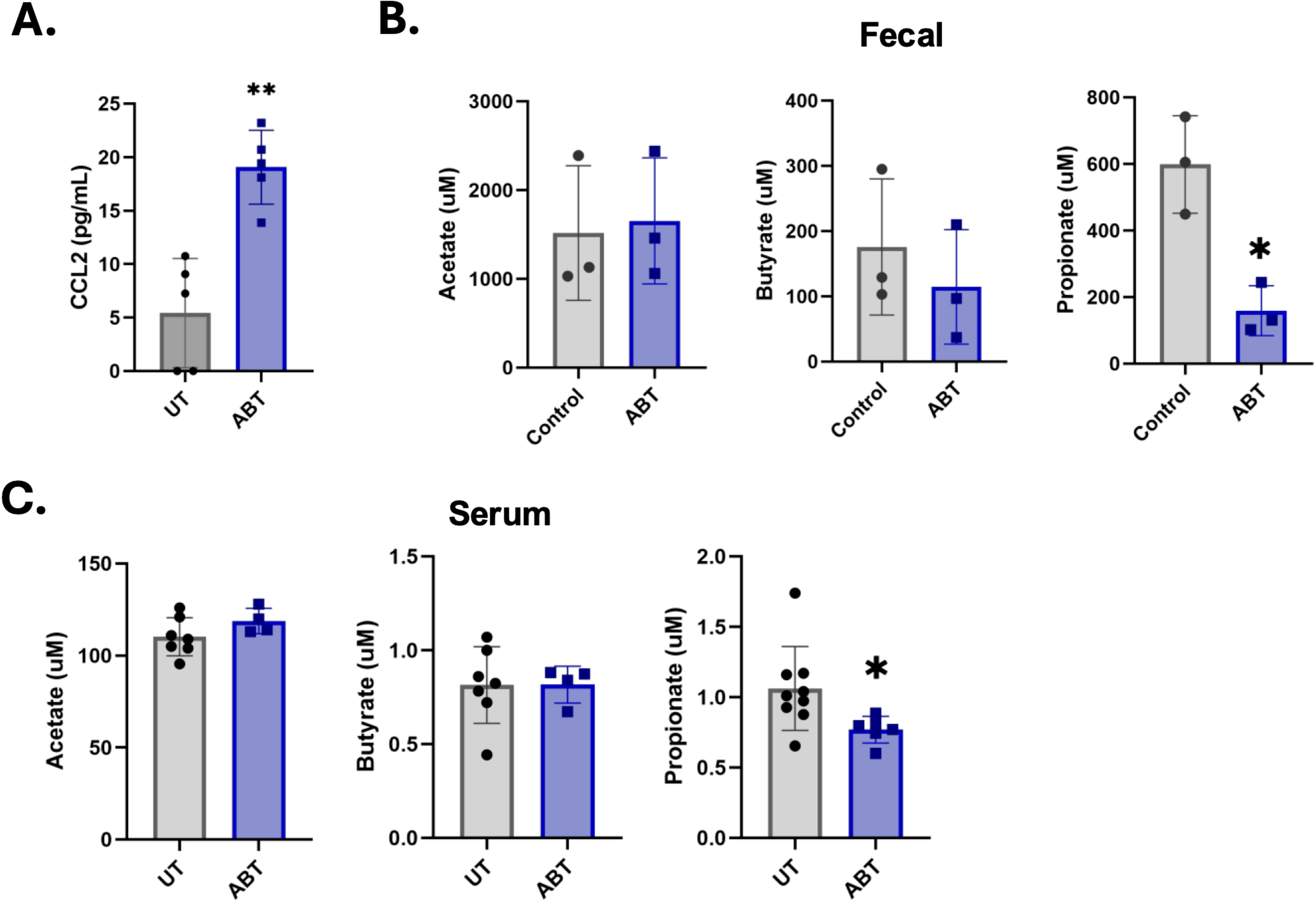
Increased CCL2 and reduced propionate in dysbiotic mice. Circulating levels of cytokines and fecal levels of SCFAs were analyzed in 8–12-week-old control (UT) and dysbiotic (ABT) female mice. A) Serum CCL2 levels measured by bead-based ELISA. B) Fecal acetate, butyrate and propionate levels. C) Serum acetate, butyrate and propionate levels. *p<0.05, **<0.01.

Because dysbiosis enhanced systemic inflammation, we then examined whether anti-inflammatory properties of the gut microbiota were altered in dysbiotic mice. The production of SCFAs is one of the main anti-inflammatory properties of the gut microbiota that acts by maintaining intestinal barrier functions, such as mucus layer thickness, and by directly inhibiting myeloid cell activation^52,12,10^. Also, *A. muciniphila* produces acetate and propionate as a by- product of mucin metabolism^19^. Therefore, we measured SCFA abundance in the fecal content and serum of dysbiotic mice. Propionate was significantly reduced in the intestinal tract and circulation of dysbiotic mice (Figure 1B-C) but other SCFAs were not affected. As such, increased systemic inflammation in dysbiotic mice is related to reduced production of propionate by the gut microbiota.

### Dysbiosis disrupts gut barrier properties

Gut barrier integrity has a profound effect on controlling immune responses distal to the gut^53^, and *A. muciniphila* is known to promote gut barrier integrity^54^. Furthermore, propionate increases mucin secretion by goblet cells^52^, promoting gut barrier function. Therefore, we next examined whether dysbiosis perturbs gut barrier properties and whether restoring *A. muciniphila* to the gut of dysbiotic mice ameliorates these effects. To examine the effects of restoring *A. muciniphila,* exogenous human *A. muciniphila* (hAKK) was administered to dysbiotic female mice immediately after cessation of antibiotics at 3 weeks of age using a voluntary oral administration protocol as previously described ^51^. Endogenous mouse *A. muciniphila* (mAKK) was absent from the stool of dysbiotic female mice as determined by qPCR (Figure 2A), which was consistent with metagenomic shotgun sequencing (Figure S1). The absolute abundance of the administered hAKK was assessed by generating a standard curve (Figure S2) and this bacterium was present at concentrations between 10^6^ to 10^7^ gene copies per gram of stool in fecal samples of dysbiotic female mice up to 8 weeks after feeding a single dose of 10^9^ CFU of *A. muciniphila* immediately after cessation of antibiotics (Figure 2B). These findings confirm that a single oral delivery of *A. muciniphila* colonizes the intestinal tract with this bacterium and can be used to examine how it affects transplant rejection.

**Figure 2.**
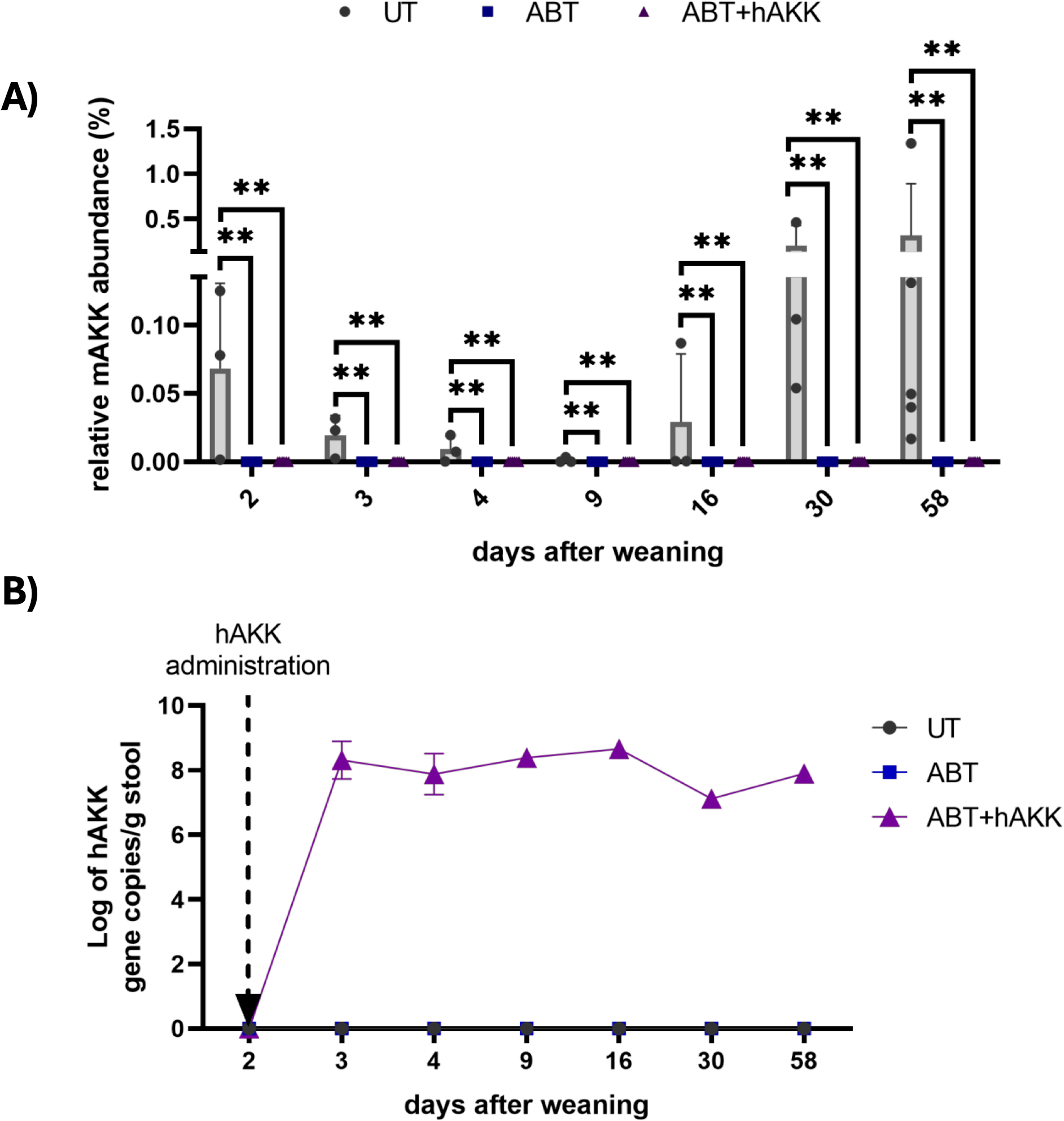
Endogenous and exogenous levels of *A. muciniphila* in fecal samples. *A. muciniphila* abundance was measured by qPCR over a period of 56 days after weaning in untreated (UT), dysbiotic (ABT), and dysbiotic mice fed exogenous *A. muciniphila* (ABT+hAKK). A) Relative abundance (%) of endogenous mouse *A. muciniphila* (mAKK) relative to pan-16S to confirm the presence or absence of mAKK between UT and ABT mice. B) Gene copies per gram of stool of exogenous human *A. muciniphila* (hAKK) was quantified to confirm the persistence of hAKK after voluntary feeding. *p<0.05, **p<0.01.

After confirming the persistence of *A. muciniphila* in the gut microbiota and its absence in dysbiotic mice, colon samples were collected and mucosal thickness as well as colon length were measured. A significant reduction in mucosal thickness was observed in the proximal colon of dysbiotic mice (Figure 3A-B). Also, colon length was significantly reduced in dysbiotic mice compared to controls (Figure 3C). Colonization of dysbiotic mice with *A. muciniphila* completely restored mucosal thickness and colon length to control levels (Figure 3B&C). Together, these data indicate that delaying colonization of the gut microbiota until after weaning perturbs gut barrier integrity and colonic development in adults, and that these changes are prevented by colonization with *A. muciniphila*.

**Figure 3.**
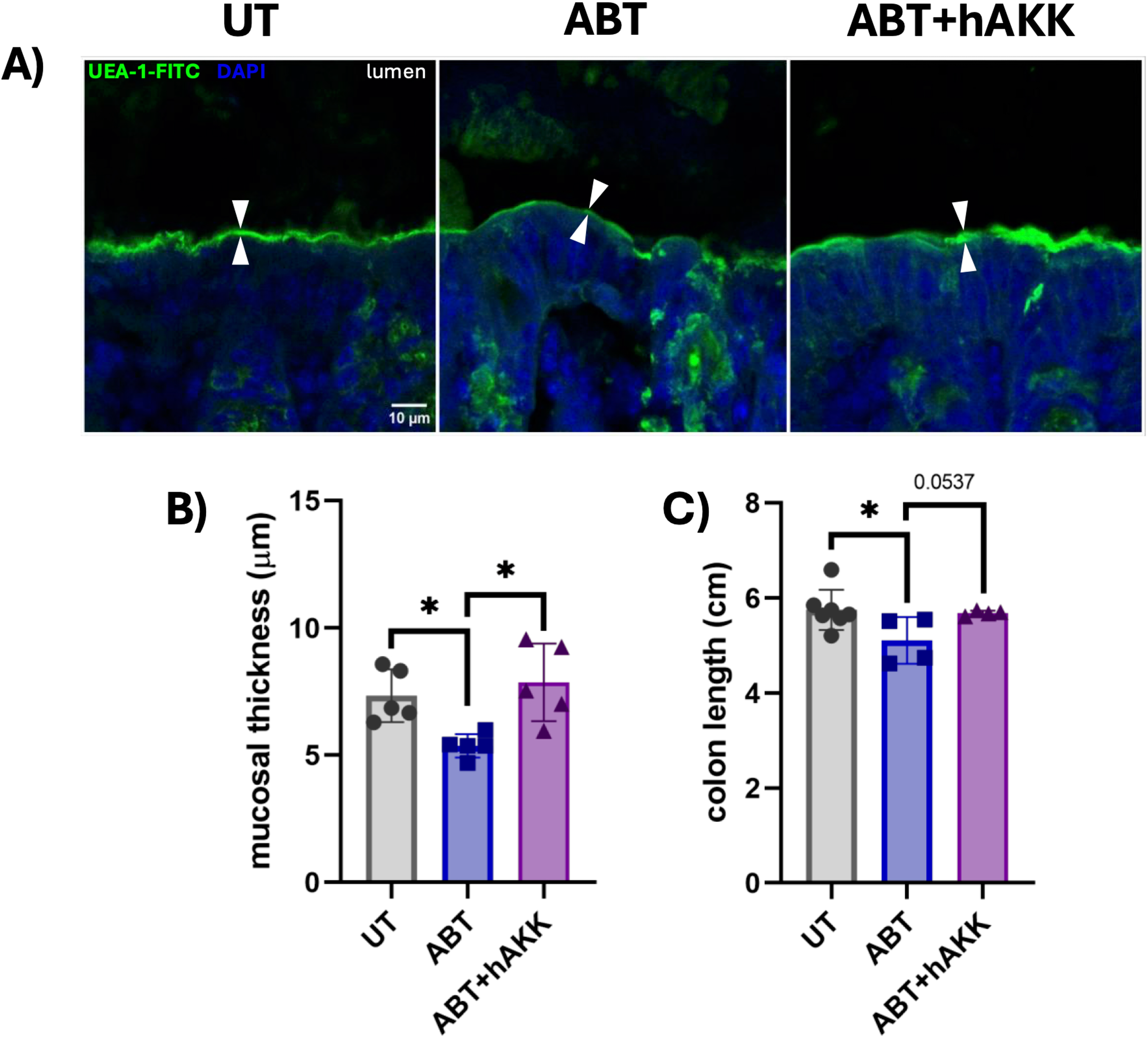
*A. muciniphila* restores gut barrier integrity in dysbiotic mice. Colon samples were collected from 8–12-week-old UT, ABT and ABT+hAKK mice. A) UEA-1-FITC staining in proximal colon. The colonic mucosa is indicated by white arrows. B) Mucosal thickness and C) Colon length were measured. *p<0.05.

### Voluntary oral administration of *A. muciniphila* ameliorates dysbiosis-induced medial injury and luminal occlusion in transplant arteries

We next examined the effect of *A. muciniphila* on dysbiosis-mediated transplant arterial injury by transplanting aortic segments from Balb/c donors into 8-12 week old C57Bl/6 recipients that had control gut microbiota, dysbiotic gut microbiota, or dysbiotic gut microbiota colonized by *A. muciniphila*. In this model, a reduction in the thickness of the medial smooth muscle cell layer can be measured as an indicator of early arterial injury reflective of acute rejection that is caused by destruction of the medial layer of arteries by myeloid and T cells^55^. Intimal thickening and luminal occlusion also develops at later time-points that is reflective of transplant arteriosclerosis, which is a main cause of late heart transplant failure ^55^. Grafts were harvested at 30 days after transplantation and cross-sections stained with H&E to assess the morphology of transplant arteries (Figure 4A). A significant reduction in medial thickness and area was observed in dysbiotic female mice (Figure 4B-C), consistent with previous findings^14,36^. Greater intimal thickening and luminal occlusion was also observed in dysbiotic mice (Figure 4D-E). Notably, colonization of dysbiotic mice with *A. muciniphila* completely ameliorated the exacerbation of medial injury and luminal occlusion (Figure 4A-E). Altogether, this shows that *A. muciniphila* in the gut microbiota inhibits acute rejection and chronic remodelling of transplant arteries caused by dysbiosis.

**Figure 4.**
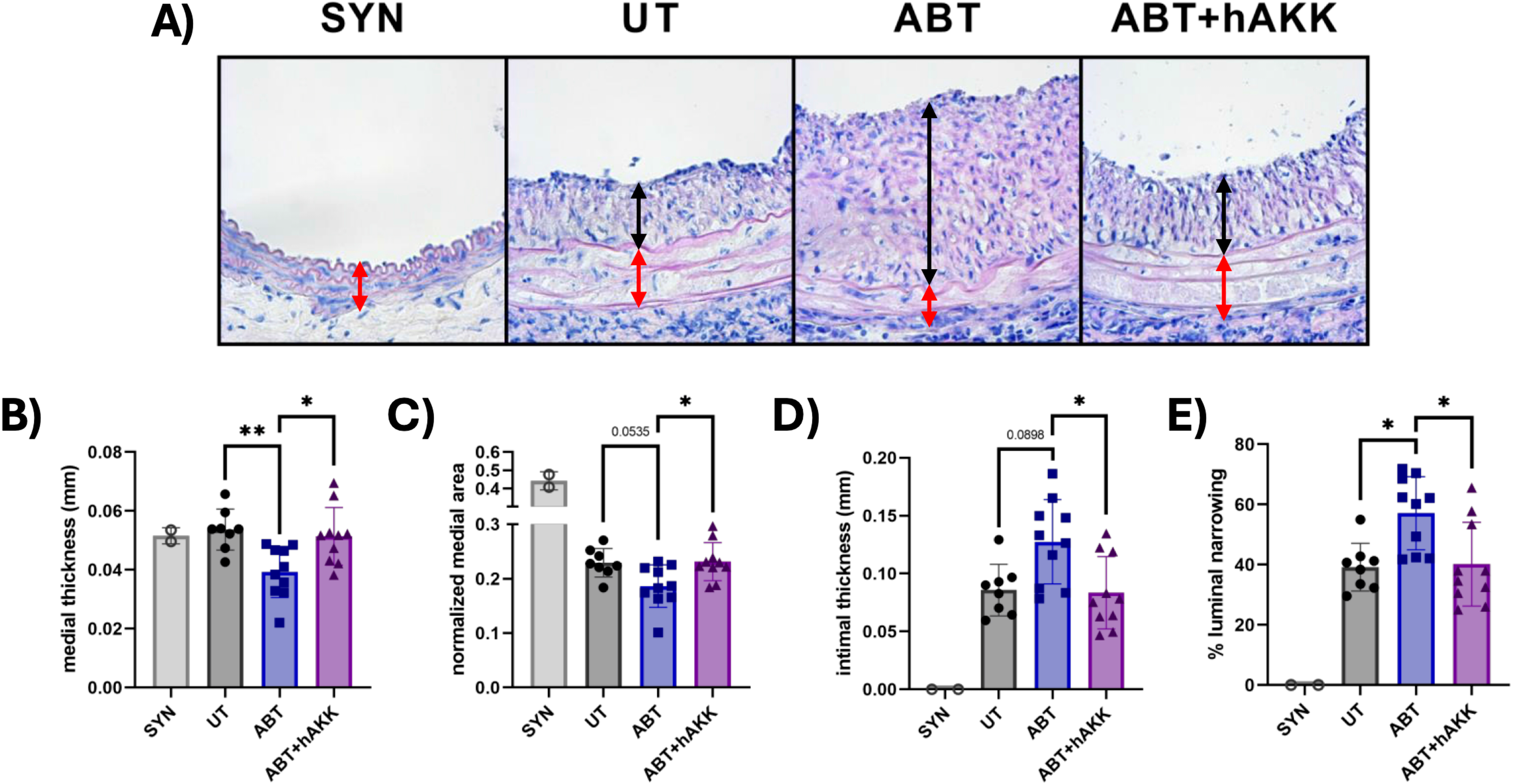
Restoring *A. muciniphila* in the gut microbiota of dysbiotic transplant recipients reduces medial thinning and luminal occlusion. Mice received arterial allografts from sex-matched recipients between 8-12 weeks of age. Syngrafts (SYN) were performed as controls. Artery grafts were harvested 30 days after transplantation and H&E stained to investigate vascular injury. A) H&E representative staining. Black bars indicate intimal growth. Red bars indicate medial thickness. B) Medial thickness, C) Normalized medial area, D) Intimal thickness, and E) % Luminal narrowing were quantified. *p<0.05, **p<0.01.

### *A. muciniphila* prevents early macrophage accumulation in allograft arteries

We next examined the effect of *A. muciniphila* on the recruitment and retention of innate immune cells to transplant arteries at day 2 post-transplantation, a time-point in which there is innate immune-mediated injury of transplants that triggers subsequent pathological changes. Macrophages were present at similar levels in the adventitia of syngrafts and allografts placed into mice with control microbiota (Figure 5A), which likely represents early infiltration in response to ischemia reperfusion injury. A significant 3.4-fold increase in macrophage accumulation was observed in the adventitia of transplant arteries from dysbiotic female recipients as compared to controls (Figure 5A). At this early time-point, no macrophages were observed in the media (Figure 5A). Notably, colonization of dysbiotic mice with *A. muciniphila* completely prevented the increase in macrophage accumulation caused by dysbiosis. Our previous findings showed that dysbiosis enhances neutrophil accumulation at day 7 post-transplantation but no differences in neutrophils were observed between any of the groups at day 2 post transplantation, indicating that early macrophage responses precede neutrophil infiltration (Figure 5B). In addition to innate immune cells, we also examined the accumulation of T cells. There was minimal accumulation of these adaptive immune cells at day 2 post-transplantation and no differences were observed between any of the groups (Figure S3). Overall, these findings reveal a regulatory role for *A. muciniphila* in inhibiting macrophage-mediated injury of transplant arteries that is caused by dysbiosis of the gut microbiota.

**Figure 5.**
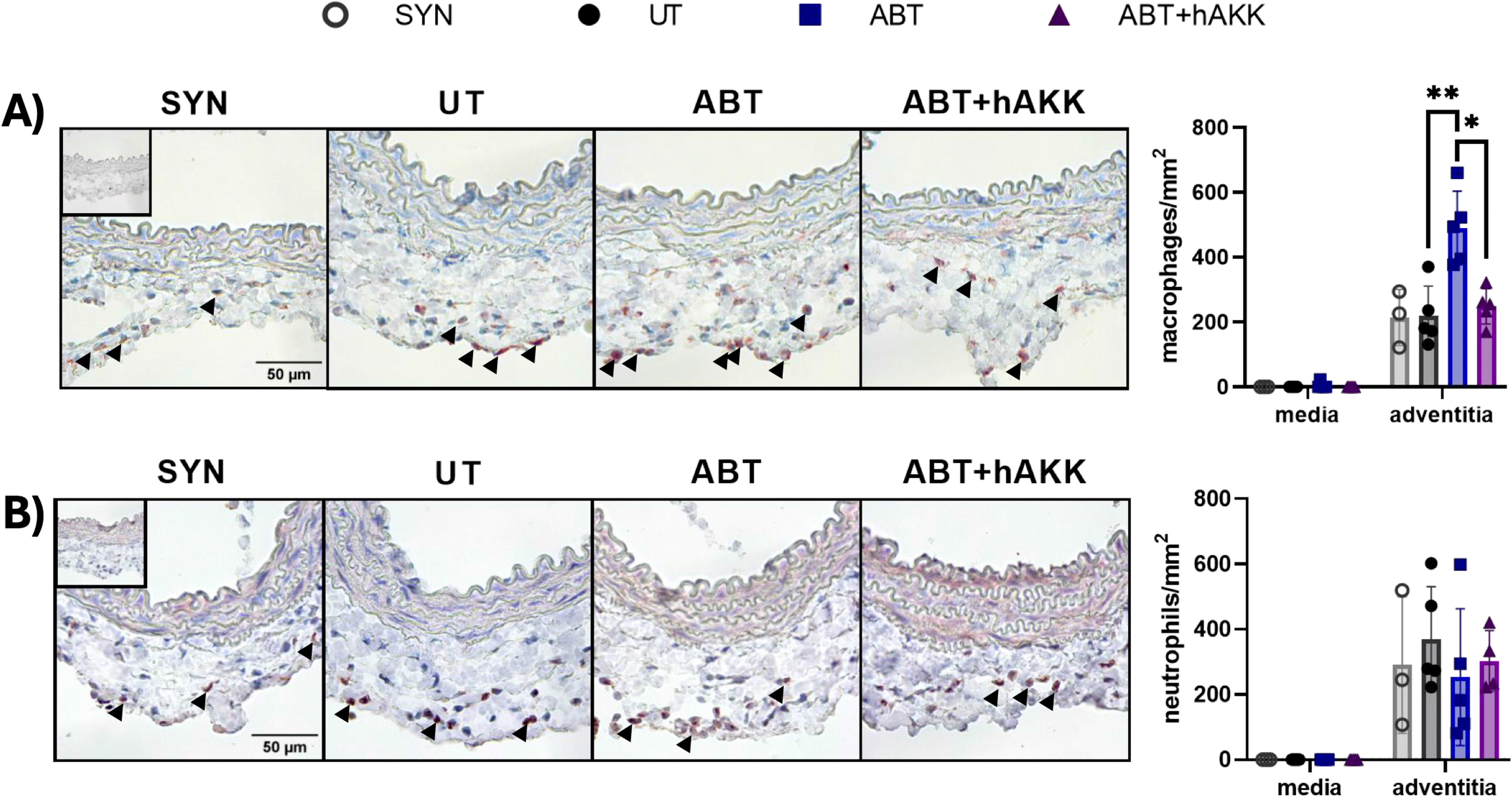
*A. muciniphila* reduces macrophage infiltration into allograft arteries of dysbiotic recipients. Artery grafts were harvested 2 days after transplantation and stained with antibodies against A) Mac-3 to visualize macrophages and B) MPO to visualize neutrophils. Graphs on the right are quantification of macrophages and neutrophils in the media and adventitia. *p<0.05, **p<0.01.

## Discussion

In our study we present new information about how early-life disruptions of the gut microbiota that delays its colonization until after weaning leads to dysbiosis that exacerbates immune-mediated injury of transplant arteries, which is involved in the acute and chronic rejection of organ transplants. Specifically, we identify that the lack of *A. muciniphila* in female murine gut microbiota leads to increased systemic inflammation and reduced intestinal barrier properties that increases myeloid responses to injure transplant arteries. Importantly, colonization of the dysbiotic gut microbiota with *A. muciniphila* completely ameliorates the exacerbation of transplant arterial injury. To our knowledge, these findings are the first to establish a protective role for *A. muciniphila* as a modifiable inhibitor of immune responses involved in acute and chronic organ transplant rejection.

Gut barrier integrity plays an important role in metabolic and cardiovascular disorders^56^. Increased gut permeability induced by diet can result in metabolic endotoxemia that contributes to vascular dysfunction and atherosclerotic lesion formation in pre-clinical models^23,57,58^. We found that early life treatment with antibiotics perturbs gut barrier integrity by reducing the thickness of the mucus layer in proximal colon. This may contribute to gut leakiness, endotoxemia and low- grade inflammation^57^. Furthermore, colon length was shortened in antibiotic treated mice, which is indicative of increased intestinal inflammation in models of colitis^59,60^. We also found that gut dysbiosis increases CCL2 expression. However, we did not investigate the source of CCL2 expression, which may be derived from the colonic epithelium or from sites distal to the gut. Indeed, CCL2 is constitutively expressed by the colonic epithelium^61^, which facilitates the recruitment of monocytes to the gut to maintain immune homeostasis.

The SCFA propionate, a by-product of mucin metabolism by *A. muciniphila*, is significantly reduced in dysbiosis caused by delaying colonization until after weaning. Production of propionate by the gut microbiota has profound effects on maintaining intestinal barrier properties and consequently immune homeostasis. Specifically, propionate can induce the production of tight junction proteins to prevent passage of microbial components into the lamina propria. This metabolite also promotes colonic epithelial cell turnover ^62,63^. We found that reduced propionate is associated with thinning of the colonic mucosa, which is consistent with other studies showing that propionate promotes mucus secretion by intestinal goblet cells^52,64^. Furthermore, propionate can help balance pro-inflammatory and tolerogenic responses within the intestine, which may also explain dysregulated CCL2 expression in dysbiotic mice with reduced fecal and serum propionate levels^65^. Propionate has also been shown to act on immune cells and directly reduce their activity^3,11,66,67^.

Graft-infiltrating macrophages are a feature of acute and chronic rejection ^68,69^. Macrophages and CCL2 are also important pathological drivers of arterial injury that causes transplant vasculopathy ^67^. We found that dysbiosis exacerbates macrophage infiltration early after transplantation and this may be facilitated by dysregulation of pathways that control CCL2 expression. This may sensitize transplant arteries to targeting by myeloid cells. Also, damage associated molecular patterns (DAMPs) are released by ischemia-reperfusion injury that induce the expression of CCL2 by donor tissue and immune cells that further recruit monocytes to allografts ^61,70^. Increased medial thinning and luminal occlusion also suggest that macrophages alter processes in both acute and chronic rejection of transplant arteries. The mechanism by which macrophages exacerbate dysbiosis-induced transplant arterial injury remain to be defined. Infiltration of macrophages contributes to tissue injury through oxidative stress by reactive oxygen species produced by NADPH oxidases and nitric oxide produced by inducible nitric oxide synthases ^71^. Their production of pro-inflammatory cytokines and chemokines may also recruit other innate immune cells, such as neutrophils, to the allograft tissue, further exacerbating immune injury^14^. Indeed, dysbiosis increases the accumulation of neutrophils after the initial increase in macrophages and neutrophil-mediated injury of transplant arteries is an early trigger that enhances the subsequent development of intimal thickening^14,72,73^.

Administering *A. muciniphila* in pre-clinical models shows promising effects in maintaining gut homeostasis and in reducing the severity of metabolic and cardiovascular disorders. Colonization of *A. muciniphila* in the gut of dysbiotic female mice prevented the effect of dysbiosis on reducing gut barrier properties. This may be mediated by propionate, which can stimulate MUC2 production in intestinal goblet cells^52^. The Amuc_1100 outer membrane protein of *A. muciniphila* also promotes gut barrier function, as it is shown to reduce endotoxemia in dysbiotic mice^26^. Supplementation of high-fat diet with *A. muciniphila* also restores metabolic health^21^. However, over-abundance of *A. muciniphila* in intestinal malignancies exacerbates inflammation and increases gut barrier permeability by over-utilizing mucus. As such, there needs to be better understanding of how the dosage and supplementation of the gut microbiota with this bacterium affects specific pathologies ^74^.

Our study provides the first evidence that the pro-health commensal bacterium *A. muciniphila* has anti-inflammatory effects that prevent arterial pathology in organ transplant rejection. This is related to the ability of *A. muciniphila* to maintain gut barrier integrity and prevent systemic inflammation. Importantly, our finding that *A. muciniphila* ameliorates macrophage-mediated transplant vascular injury is one of the first to establish its role as a regulator of immune responses in organ transplant rejection.

## Acknowledgements

The study was funded by the Heart and Stroke Foundation of Canada (J.C.C) and the Canadian Institutes of Health Research (L.E.D).

## Author contributions

JMD designed the study, performed experiments and analyzed the data, and wrote the manuscript; KR performed experiments and analyzed the data; PM, EW, WE, and CRH performed the experiments; LED analyzed the data; JCC designed the study, analyzed the data, and wrote the manuscript.

## Notes

### Competing Interest Statement

The authors have declared no competing interest.

